# Discovery of Chemical Tools for Polysorbate-Degradative Enzyme Control in Biopharmaceutical Upstream Process via Multi-Omic Profiling of Host Cell Clones

**DOI:** 10.1101/2025.01.24.634759

**Authors:** Taku Tsukidate, Ansuman Sahoo, Geetanjali Pendyala, Rong-Sheng Yang, Jonathan Welch, Sri Madabhushi, Xuanwen Li

**Author notes:** email: X.L.; S.M. These authors contributed equally: Taku Tsukidate, Ansuman Sahoo.

## Abstract

Host cell proteins are process-related impurities in biotherapeutics and can potentially pose risks to patient safety and product quality. Specifically, certain host cell–derived enzymes, including lipases, can degrade the formulation excipient polysorbate (PS) in biopharmaceutical formulations, affecting drug product stability in liquid formulations. We leveraged multi-omics approaches, including transcriptomics, proteomics, and activity-based protein profiling (ABPP), to identify mechanisms that regulate PS-degradative enzyme (PSDE) abundance and to develop strategies for their control. Comparative multiomics analysis of two monoclonal antibodies (mAb)-producing host cell clones revealed differential lipase profiles at the mRNA, protein, and enzyme activity levels and associated increased lipase activity with upregulated lipid catabolic pathways such as the fatty acid beta oxidation pathway. Further, for the first time in the literature, we identified peroxisome proliferator–activated receptor γ (PPARγ) as a key regulator of PSDEs in manufacturing Chinese Hamster Ovary (CHO) cells. Downregulation of the PPARγ pathway with its antagonists resulted in selective reduction of PSDE levels and improved PS stability without compromising mAb productivity or quality. This study highlights the potential of PPARγ modulators as chemical tools for PSDE control at the gene regulation level, offering significant implications for biopharmaceutical process development and control.

## Introduction

Host cell proteins (HCPs) are process-related impurities in biotherapeutics that originate from host cells. Regulatory agencies, such as Food and Drug Administration (FDA) and European Medicines Agency (EMA), expect that HCP impurities are well-controlled and understood and that these impurities are not a risk to patient safety or product efficacy. In addition, several HCPs are considered especially high risk of affecting product quality, stability, and/or patient safety and require control strategies for removal and monitoring.^1^ For example, polysorbates (PS) as non-ionic surfactants are widely used excipents in biophamaceutic formulations to protect proteins against interfacial stress. Over 86 % (112/126) of commercially approved antibodies by Feb 2021 have PS- 20 or −80 in their drug products.^2^ Sub-ppm levels of certain residual HCPs can degrade PS significantly enough to cause particle formation and affect product quality, which has become an industry-wide challenge especially with increased cell culture density and higher protein concentration formulation.^3,4^ We and others have identified and characterized these PS-degradative enzymes (PSDEs) many of which are lipases such as APT1^5^, CES (carboxylesterase) homologs^6^, LIPA^7,8^, LPL^7,8^, PLA2G15^9^, PLA2G7^10^, PPT1^7^, and SIAE^11^.

Achieving sufficient removal of HCPs through purification can be a challenging task in biopharmaceutical process development.^12,13^ In this regard, recent studies have evaluated various strategies to reduce the abundance of HCPs at the cell culture stage and improve the efficiency of purification processes. For example, components in cell culture media can alter the HCP profile. In one study, higher concentrations of insulin, folic acid, glycine or riboflavin decreased total HCP concentrations while the addition of magnesium chloride increased the total HCP concentration.^14^ Similarly, bioreactor process parameters such as reactor type^15^, temperature^16–18^, pH^18^, and harvest viability^19–21^, can contribute to total HCP concentration and individual HCP abundance.^22^ On the other hand, changes in media composition or process parameters tend to broadly impact cell culture performance and would require a comprehensive mechanistic understanding to enable selective reduction of HCP levels without compromising other critical quality attributes. In contrast, cell line engineering can target specific HCPs for reduction. For example, a recent study reported a substantial reduction in total HCP concentration while maintaining high productivity by knocking out several genes encoding abundant HCPs.^23^ Similarly, others demonstrated that PSDE-knockout^24^ or -knockdown^25^ cell lines reduce PS degradation. However, cell line engineering has limited applicability to pipeline programmes that have passed the cell line development stage, especially in current biopharmaceutical merger and acquisition environment.

Omics technologies such as genomics, transcriptomics, proteomics and metabolomics have advanced bioprocess understanding over the past decade.^26–28^ In particular, integration of different omics types *i.e.*, multi-omics^29^ is a powerful approach for understanding the flow of biological information from causal factors such as genetics, process parameters, *etc.* to functional and/or phenotypic consequences such as productivity, product quality profile, *etc.* For example, a recent study evaluated the impact of culture pH on quality profile, employing transcriptomics, proteomics, and metabolomics.^30^ The authors observed an association among culture pH, product glycosylation, and ER stress response pathways in multiple omics layers and speculated how culture pH would affect ER function and, in turn, product quality. Similarly, another multi-omics study evaluated the impact of cysteine feed on productivity and revealed stress responses and metabolic shifts that are associated with productivity decline.^31,32^ Nevertheless, it can be challenging to translate observational insights from omics studies into actions in bioprocess development.^28^

Here, we describe a multi-omics–guided discovery of PS control tools for upstream cell culture process. We combine transcriptomics, abundance-based proteomics, and activity-based proteomics to hypothesise mechanisms regulating PSDE abundances and identify peroxisome proliferator–activated receptor γ (PPARγ) as a key regulator. We also demonstrate that the downregulation of the PPARγ pathway significantly lowers the PLA2G7 level and reduces PS degradation.

## Results

### Lipase activity difference between clones

We cultured the two most productive clones of a proprietary monoclonal immunoglobulin G (IgG) in a typical 14-day fed-batch process in shake flasks. Clone A cells had higher cumulative cell densities (Fig. 1a) and consumed more gluatamine and produced more ammonia (Supplementary Fig. 1a–d) than Clone B cells. Despite these differences in their growth profiles, these clones had comparable titer (Fig. 1b) and product quality (monomer: 99 %; charge variants: Supplementary Fig. 1e) of IgG in their harvested cell culture fluids (HCCFs). However, the total lipase activity was significantly higher in the Clone A HCCF than in the Clone B HCCF from the seventh day of culture (Fig. 1c). The total lipase activity in the Clone A HCCF increased as the culture matured while its counterpart in the Clone B HCCF remained largely unchanged, culminating in an approximately two-fold difference at day 14. These results suggest that HCCFs from equally productive clones can significantly differ in lipase profile and that clone selection may have implications for downstream PS degradation control strategy.

**Fig. 1.**
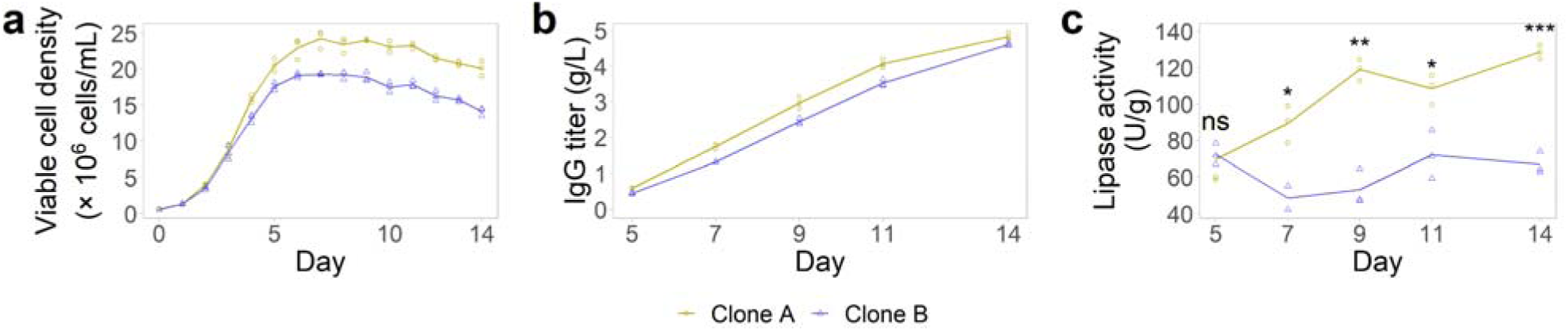
Fed-batch culture profiles. **a** viable cell density, **b** IgG titer, and **c** total lipase activity per titer. Student’s *t*-test. ns: not significant; *: p < 0.05; **: p < 0.01; ***: p < 0.001. n_Clone A_ = 3. n_Clone B_ = 3.

### Identification of differentially abundant and/or active lipases in HCCF

To identify differentially abundant lipases between the Clone A HCCF and the Clone B HCCF, we performed a data-independent acquisition (DIA)– based quantitative proteomics analysis on the fifth, seventh, nineth, eleventh, and fourteenth days of culture.^33^ Overall, we quantified 1961 HCPs across 30 samples with three biological replicates. Several lipases were significantly more abundant in the Clone A HCCF than in the Clone B HCCF, including known PSDEs such as LIPA, PLA2G15, PLA2G7, and PPT1 (Fig. 2a, Supplementary Fig. 2). Among these, PLA2G7 was consistently at least twice as abundant at all time points, whereas LIPA, PLA2G15 and PPT1 became increasingly more abundant from the seventh day, miroring the total lipase activity trend (Fig. 2b). On the other hand, the LPL and SIAE levels were similar between the clones. In addition, the Clone A HCCF contained higher levels of cathepsins whereas the Clone B HCCF contained higher levels of a matrix metalloproteinase and a protein disulfide isomerase as other high-risk HCPs (Supplementary Fig. 2). However, abundance measurements alone cannot probe the functional states of enzymes. To identify active lipases in the HCCFs, we performed an activity-based protein profiling (ABPP) of these samples employing the active site–directed chemical reporter FP-biotin^10,34^ (Fig. 2c, d, Supplementary Fig. 3). PLA2G7 was more robustly enriched from the Clone A HCCF than from the Clone B HCCF, perhaps reflecting the difference in abundance. In contrast, LIPA was not significantly enriched from either HCCFs despite being present in both, which is consistent with its acidic pH optimum.^35–37^ Overall, more lipases were active in the Clone A than in Clone B (Fig 2e). These results indicate that the Clone A HCCF indeed contains higher levels of certain active lipases such as PLA2G7, PPT1 and, to somewhat lesser extent, PLA2G15 than the Clone B HCCF.

**Fig. 2.**
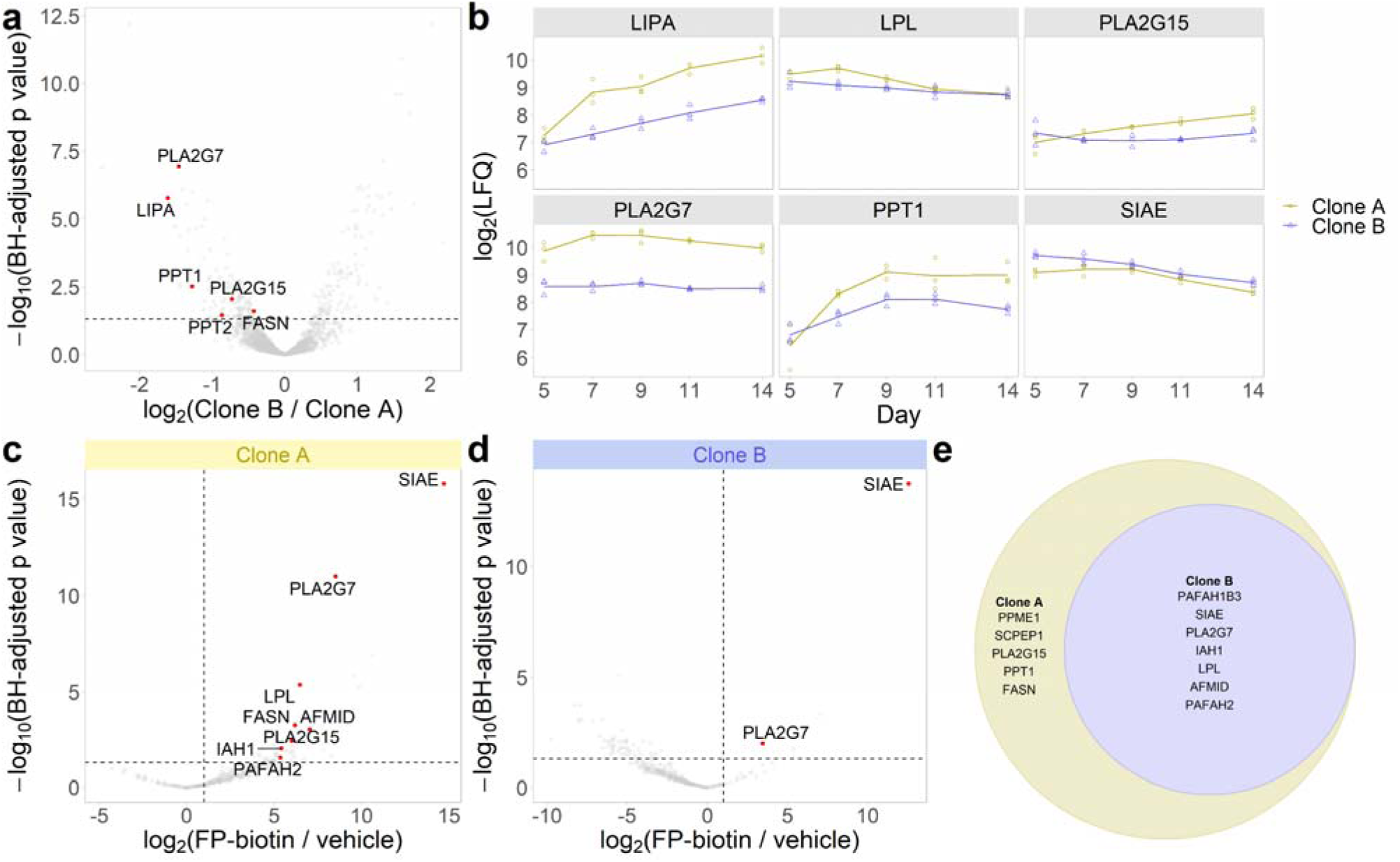
LC-MS/MS–based analysis of lipase abundance and activity in HCCF. **a** Comparison of HCP abundance between Clone A and Clone B at day 14. The horizontal line indicates adjusted p value < 0.05 in empirical Bayes–moderated *t*-test. Lipases with p < 0.05 are highlighted. n_Clone A_ = 3. n_Clone B_ = 3. **B** Time-course profiles of known PSDEs in HCCF. ABPP of lipases at day 14 in Clone A (**c**) and in Clone B (**d**). The vertical line indicates log_2_(FP-biotin / vehicle) > 1 and the horizontal line indicates adjusted p value < 0.05 in empirical Bayes–moderated *t*-test. Lipases with p < 0.05 are highlighted. n_FP-biotin_ = 3. n_vehicle_ = 3. **e** Comparison of active lipases between Clone A and Clone B. Active lipases were combined across all time points.

### Correlation analysis of transcriptome and proteome in cells

Various mechanisms regulate HCP levels beyond transcription. Thus, combining a transcriptome-level analysis and a proteome-level analysis would lead to better understanding of these regulatory mechanisms. On one hand, we previously performed a bulk RNA-seq analysis of Clone A cells and Clone B cells cultured in fed-batch bioreactors operated under various conditions.^21^ We leveraged this existing dataset by extracting a subset of data where culture conditions closely resembled current shake-flask conditions. This subset included the quantification of 12,105 mRNAs in 11 bioreactors at six timepoints. On the other hand, we performed a DIA–based quantitative proteomics analysis of cell pellets from current shake-flask runs at five timepoints and quantified 6103 proteins across 30 samples. With these related transcriptomics and proteomics datasets in hand, we analysed Spearman pair-wise correlations of all mRNA–protein pairs (n = 4953) by matching clones and timepoints (n = 6) (Fig. 3a). The median correlation coefficient for timepoint-wise analysis was 0.09, whereas the median correlation coefficient for cumulative analysis *i.e.*, correlation between cumulative sum of mRNA and protein was 0.66. This suggests that the cumulative mRNA level may serve as a better proxy for protein levels. To evaluate the functional implications of divergence between mRNA and protein levels, we performed a Gene Set Enrichment Analysis (GSEA) using a gene list ranked by the Spearman correlation coefficient. Highly correlated pairs belonged to energy and lipid metabolism processes whereas poorly correlated pairs belonged to housekeeping functions such as ribosomal and spliceosomal processes (Fig. 3b–d). Previous studies reported similarly poor correlations between mRNA and protein levels of these housekeeping genes.^38,39^ Known PSDEs tend to be involved in the lipid catabolic process and their cumulative mRNA levels well correlated with their protein levels (Fig. 3e), while spliceosomal components such as SRSF1 were among the worst correlating pairs (Fig. 3f). These results suggest that the combination of transcriptomics and proteomics should boost confidence in hypothesizing potential mechanisms regulating PSDE levels.

**Fig. 3.**
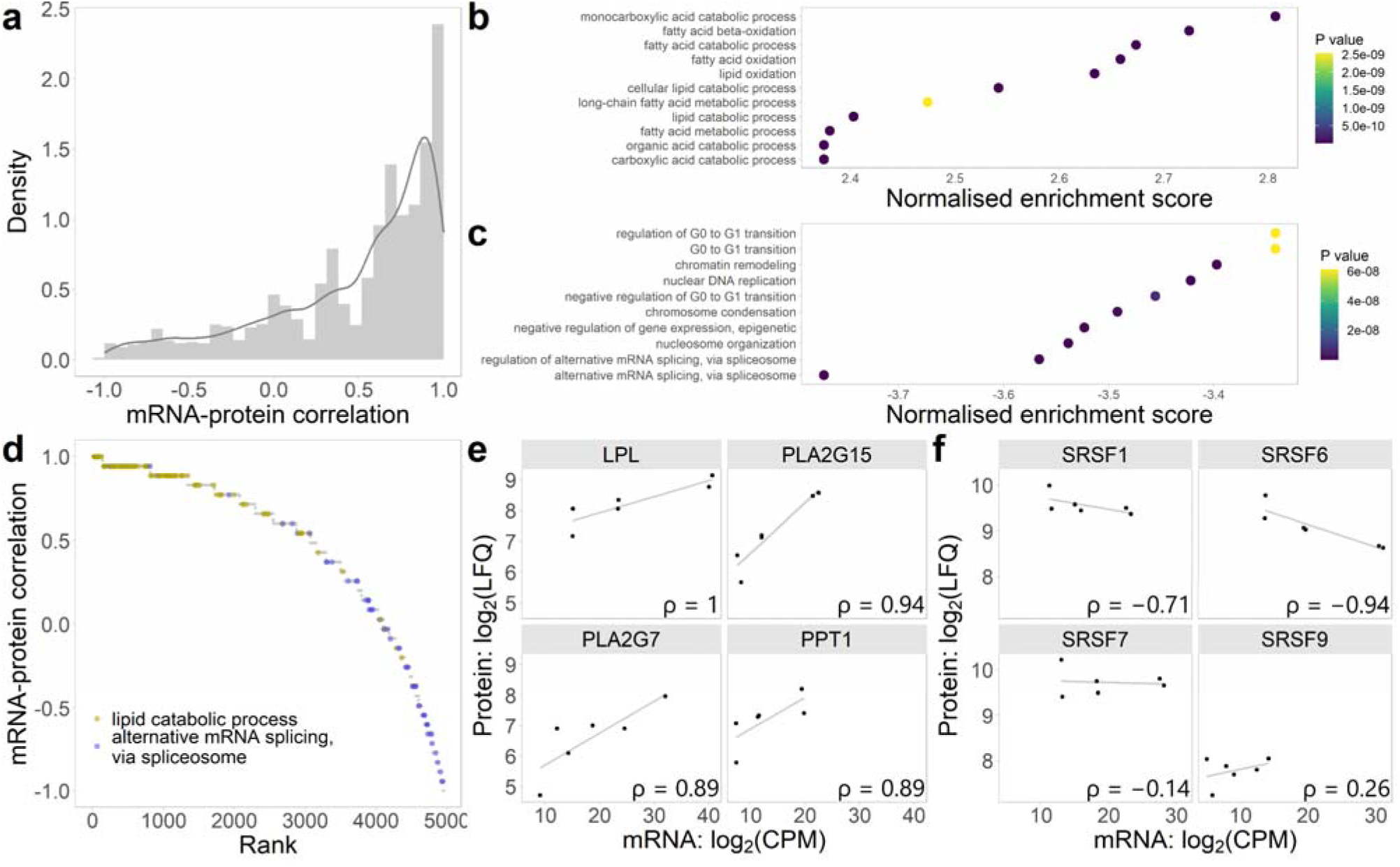
Synergy and discordance between mRNA and protein levels in cells. **a** Distribution of Spearman correlation coefficients between cumulative mRNA levels and protein levels. GSEA of the mRNA–protein level correlations for Gene Ontology Bioprocess (GO BP) terms, showing the most correlated pathways (**b**) and the least correlated pathways (**c**). **d** Ranked mRNA–protein level correlations with highlighted markers for lipid catabolic process (yellow) and alternative mRNA splicing, via spliceosome (blue). mRNA levels versus protein levels for select PSDEs (**e**) and select spliceosome components (**f**). ρ: Spearman’s correlation coefficient.

### Identification of pathways associated with high PSDE levels

Next, we leveraged these datasets to derive a hypothesis for a mechanism regulating PSDE levels in CHO cell bioprocess. A hierarchical clustering analysis of the cell pellet proteomics data (6103 proteins × 30 samples) stratified the samples into clusters characterized by cell culture phase *i.e.*, early stationary phase *versus* late stationary and decline phase and by clone (Fig. 4a). Consistent with the HCCF, PLA2G7 was more abundant in Clone A than in Clone B and LPL and PLA2G15 were similarly abundant (Fig. 4b). In contrast, intracellular levels of LIPA and PPT1, both of which were significantly more abundant in the Clone A HCCF than in the Clone B HCCF, were similar between the clones, suggesting that the abundances of these two PSDEs in HCCF may be differentially impacted by post-translational mechanisms such as secretion. Indeed, the mRNA levels of the PSDEs trended similarly to their cellular protein levels (Supplementary Fig. 4a). To further investigate mechanisms that regulate PSDE expression levels and the lipase activity, we performed a differential abundance analysis between the two clones at the seventh day, when significant difference in the total lipase activity emerged in HCCFs (Fig.1c and Fig. 4c). Overall, 224 proteins were significantly more abundant in Clone A whereas 1732 proteins were significantly more abundant in Clone B (adjusted p value < 0.01). Interestingly, a GSEA indicated that proteins involved in fatty acid beta-oxidation and other lipid catabolic pathways were significantly enriched in Clone A (Fig. 4d, Supplementary Fig. 5). For example, several acyl-CoA dehydrogenases such as ACAD10^40^ and ACADVL^41^ that catalyzes the first step of fatty acid beta-oxidation were significantly more abundant in Clone A. Concurrently, a GSEA using the transcriptomics data supported the enrichment of fatty acid beta-oxidation in Clone A (Fig. 4e). For example, *Acaa2,* which catalyzes the final step of fatty acid beta-oxidation^42^, was significantly more abundantly expressed in Clone A (Supplementary Fig. 4b). Similarly, *Pparg*, which modulates the expression of genes involved in lipid metabolism and controls fatty acid beta-oxidation^43^, was also among the 122 genes that were significantly upregulated in Clone A (Supplementary Fig. 4b). In contrast, proteins involved in chromatin remodeling and spliceosomal processes were enriched in Clone B, which was not corroborated at the mRNA level (Supplementary Fig. 4c, d). These results suggest that the upregulation of lipid catabolic pathways including fatty acid beta-oxidation is associated with increased PSDE levels and lipase activity.

**Fig. 4.**
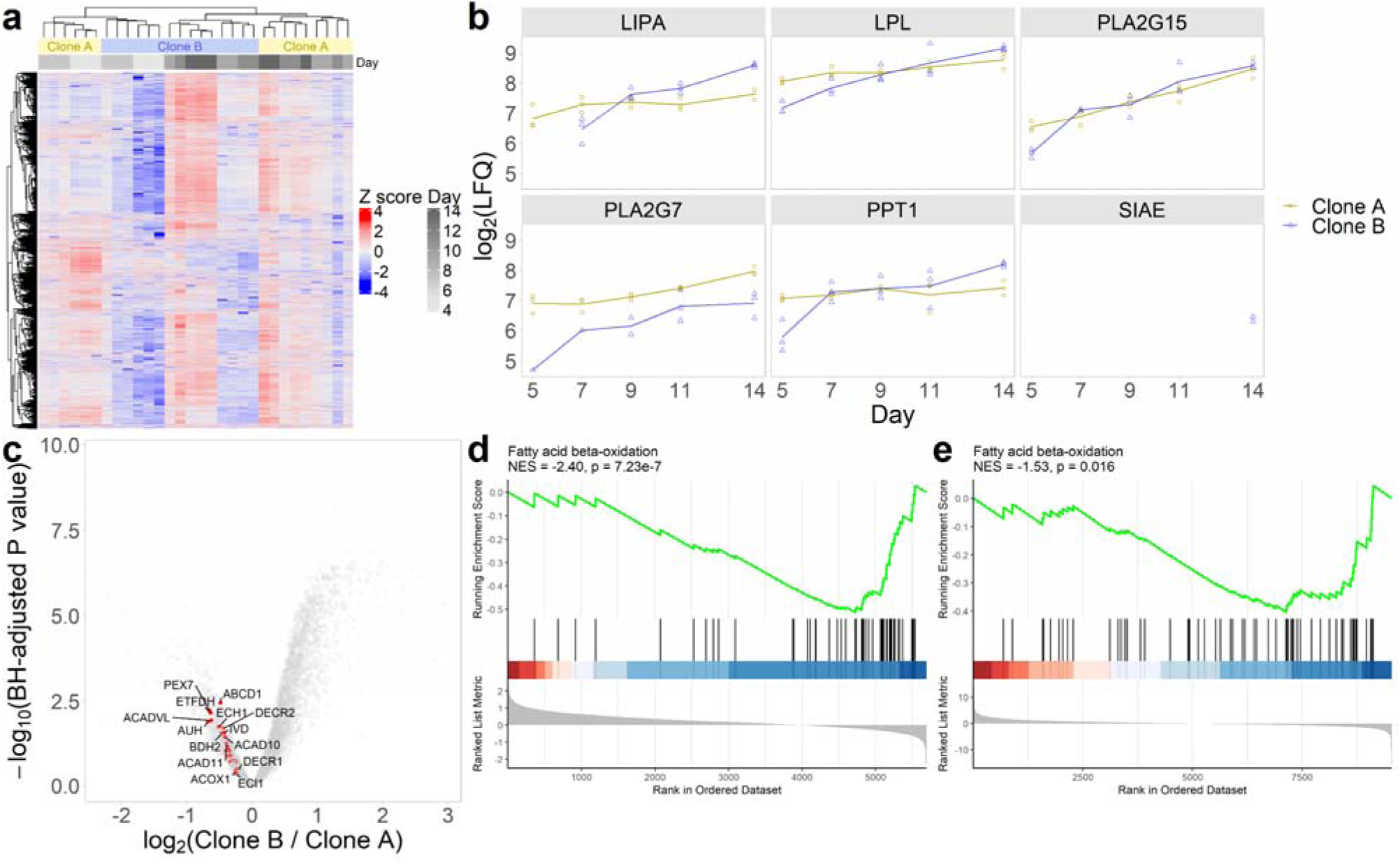
LC-MS/MS–based analysis of protein abundance in cells. **a** Heatmap visualization of 6103 quantified proteins in Clone A and Clone B cells across five different time points. **b** Time-course profiles of known PSDE protein abundances in cells. **c** Comparison of protein abundances between Clone A and Clone B at day 7 with highlighted protein markers for fatty acid beta-oxidation leading-edge. Empirical Bayes–moderated *t*-test. n_Clone A_ = 3. n_Clone B_ = 3. GSEA of differentially expressed genes between Clone A and Clone B using the proteomics data (**d**) or the transcriptomics data (**e**), showing significant enrichment of fatty acid beta–oxidation in Clone A.

### Selective reduction of PLA2G7 level with PPARγ antagonists

Peroxisome proliferator–activated receptor γ (PPARγ) is a nuclear receptor that functions as a transcription factor and regulates the expression of genes involved in lipid metabolism and beyond.^43^ While early studies characterized PPARγ as a crucial regulator of adipogenesis and as a target of a class of insulin-sensitizing therapeutics called thioglitazones, more recent studies have revealed its diverse roles in other cell types. For example, PPARγ induces anti-inflammatory phenotype in macrophages through several mechanisms including the promotion of fatty acid oxidation and glutaminolysis.^44–46^ The enrichment of lipid metabolic pathways and the higher glutamine consumption in Clone A suggest an upregulation of PPARγ-regulated pathways in this clone. Indeed, PPARγ-regulated genes^47^ were significantly over-represented among genes that were upregulated in Clone A both at the mRNA level (Fisher’s exact test, p = 0.0019) and the protein level (Fisher’s exact test, p = 0.044). Moreover, several studies have demonstrated that PPARγ ligands can modulate *PLA2G7* expression levels in monocytic cell line models.^48,49^ Thus, we hypothesized that downregulation of these pathways would result in lower PSDE levels. To test this hypothesis, we cultured Clone A in a batch mode in the absence and the presence of the non-covalent PPARγ antagonist SR1664^50^ (1 μM at days zero and two), which directly binds PPARγ in cells^51^ and inhibits the transcriptional function. SR1664 did not significantly affect cell growth, cell viability or IgG titer (Fig. 5a–c). However, the SR1664 treatment indeed led to significant decreases by ≥ 20 % in the protein levels of the PPARγ-regulated genes *Fabp4*, *Lrp1* and, importantly, *Pla2g7* as well as a decreasing trend in the LPL protein level in the HCCF (Fig. 5d). To evaluate potential proteome-wide impact of PPARγ antagonism, we performed a DIA–based quantitative proteomics (Fig. 5e) and quantified 2717 HCPs. Overall, 41 proteins were significantly more abundant in the control group whereas 14 proteins were significantly more abundant in the SR1664 group (Supplementary Table 1). Besides PLA2G7, the three high-risk HCPs CTSO^52,53^, MMP19^54^ and TGFB1^55^ and the PSDE SIAE were significantly less abundant in the SR1664 group. Moreover, a GSEA indicated significant depletion of proteins involved in lipid catabolism (Fig. 5f, g), suggesting that PPARγ is indeed an important regulator of these processes in CHO cells. On the other hand, we did not observe significant increase in other lipase or high-risk HCP levels in the SR1664 group. In addition, a treatment with an analogous PPARγ inverse agonist^56^ similarly decreased the PLA2G7 level (Supplementary Fig. 6, Supplementary Table 2), suggesting the on-target effect of PPARγ modulation.

**Fig. 5.**
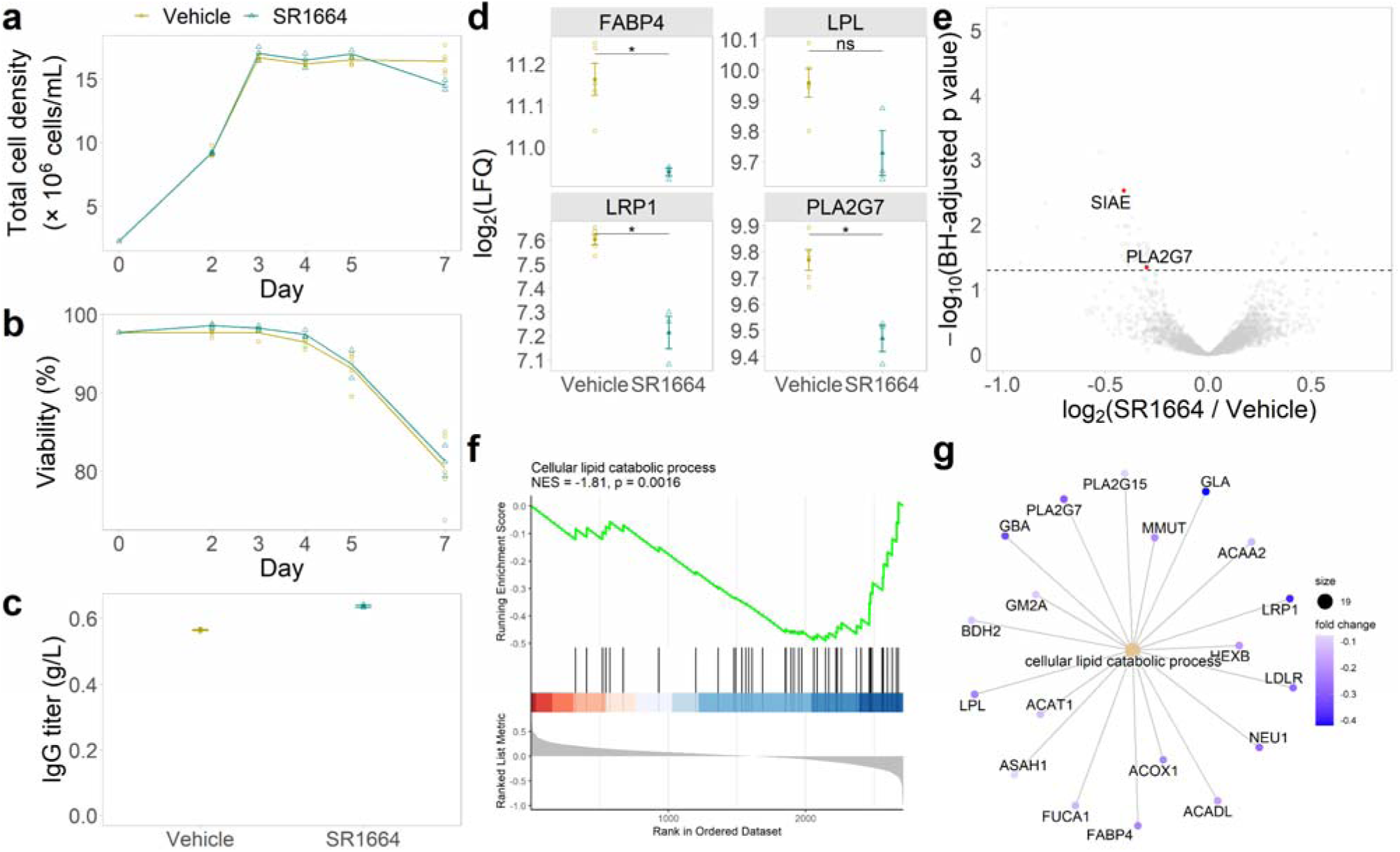
Selective reduction of PSDE levels in HCCF with PPARγ antagonist. SR1664 did not significantly affect cell growth (**a**), cell viability (**b**), or IgG titer (**c**). **d** Protein levels of select PPARγ- regulated genes. Error bars represent the standard errors of mean. Student’s *t*-test. ns: not significant; *: p < 0.05. **e** Comparison of HCP levels between vehicle-control and SR1664-treatment groups, showing selective reduction of PLA2G7 and SIAE. The horizontal line indicates adjusted p value < 0.05 in empirical Bayes–moderated *t*-test. GSEA of differentially abundant proteins between vehicle-control and SR1664-treatment groups, showing significant depletion of cellular lipid catabolic process (**f**) and fold changes of its associated proteins (**g**). n_Vehicle_ = 6. n_SR1664_ = 3.

### Improvement of polysorbate stability in Protein A pool

Encouraged by these results, we assessed whether SR1664 could serve as a chemical tool to control PS degradation. We cultured Clone A in the fed-batch bioreactor in the absence or presence of SR1664 (1 × 10^-15^ mol/cell) for 12 days. SR1664 did not significantly affect either cell viability (Fig. 6a) or IgG titer (Fig. 6b) but reduced total lipase activity in the HCCF approximately by half (Fig. 6c). In addition, the SR1664 treatment group produced less dense cultures and consumed less glutamine than the vehicle control group, which mirrors the difference between the less lipolitic Clone B and the more lipolitic Clone A (Supplementary Fig. S7 and S1). To evaluate the impact of the lipase downregulation on downstream purification, we performed protein A affinity chromatography and analyzed these protein A pool (PAP) samples with respect to lipase activity and product quality. Purification yield ranged 90–95 % and were comparable between the two groups. Importantly, SR1664 did not significantly alter the charge variant profile (Supplementary Fig. S7g). Gratifyingly, total lipase activity was 27 % lower in the SR1664 treatment group (Fig. 6d), which translated to 34 % improvement of PS-80 stability over two weeks (Fig. 6e). These results demonstrate that SR1664 can reduce the PS-degrading potential without compromising productivity or product quality.

**Fig. 6.**
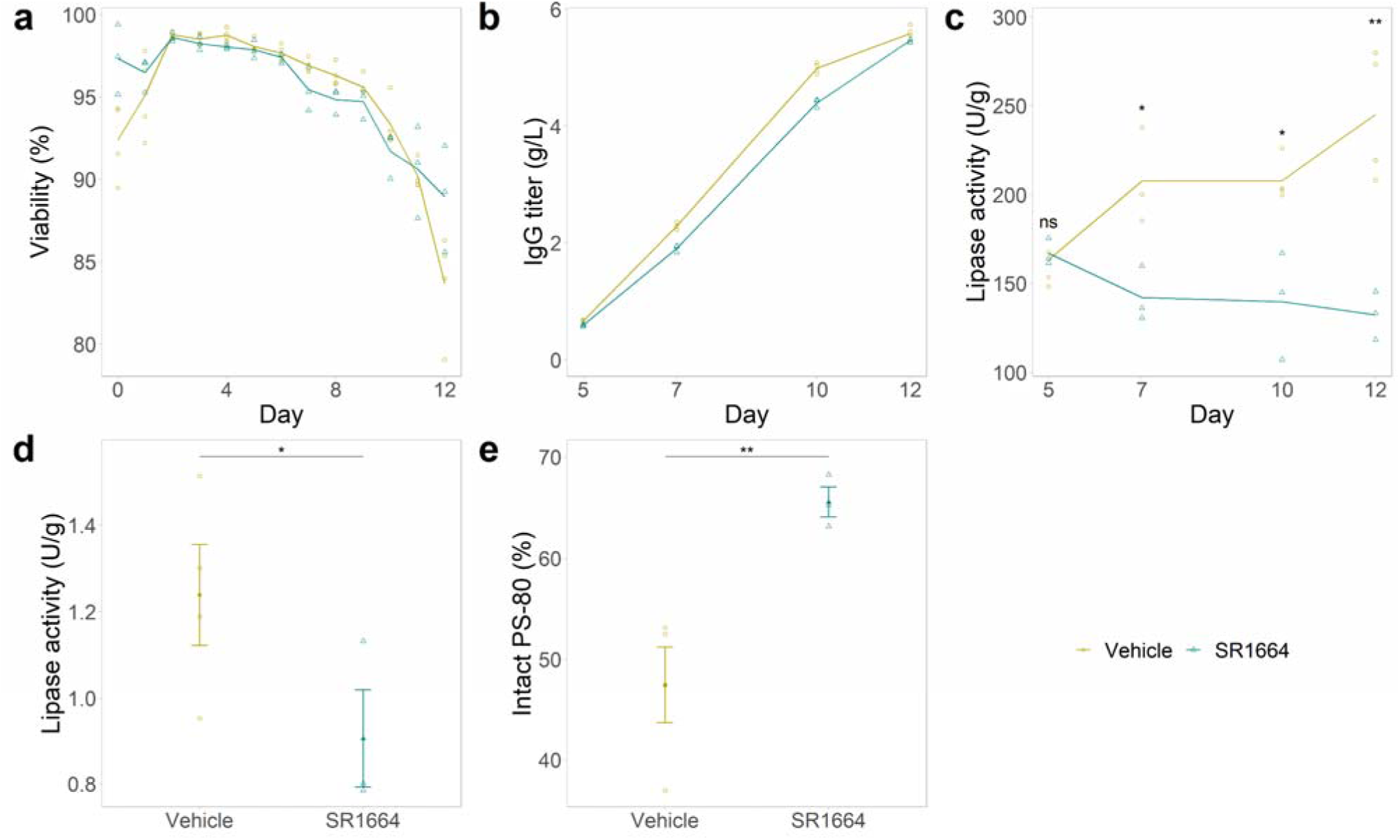
Improvement of PS stability with PPARγ antagonist. SR1664 did not significantly affect cell viability (**a**) or IgG titer (**b**) in fed-batch culture. Total lipase activity per titer in HCCF (**c**) and PAP (**d**). **e** Intact PS-80 level in PAP after two-week incubation at 37 °C expressed as a percentage of its level before incubation. Error bars represent the standard error of mean. Student’s *t*-test. ns: not significant; *: p < 0.05; **: p < 0.01. n_Vehicle_ = 4. n_SR1664_ = 3.

## Discussion

HCP control is an essential part of biopharmaceutical process development for ensuring patient safety and product quality. However, achieving sufficient removal of HCPs through purification is a challenging task and can be a bottleneck in biopharmaceutical production.^12,13^ To alleviate these challenges, recent studies have explored alternative strategies to reduce the loads of HCPs through cell culture medium composition, bioreactor process parameters, or cell line engineering with varying degrees of success. In this regard, systems biology approaches including omics data analysis can enhance these efforts.^28^ Secretory pathway modelling has identified several HCPs that would liberate a large portion of cellular resources upon deletion, and a multi-plexed knockout of these HCPs has indeed resulted in higher mAb productivity and purity.^23,57^

Here, we leveraged multi-omics data analysis to derive actionable insights into regulatory mechanisms of PSDE levels. We focused on two clones that were similar in terms of titer and quality of IgG. Comparative profiling at the transcriptome and the proteome levels revealed the association between the increased PSDE levels and the upregulation of cellular lipid catabolic pathways such as fatty acid beta oxidation. Accordingly, we hypothesized that downregulating these pathways with PPARγ modulators would lower PSDE levels. Indeed, the PPARγ antagonist SR1664 reduced the PLA2G7 level in the HCCF and improved PS stability after purification. Importantly, SR1664 diminished neither cell viability, productivity, nor quality of IgG, which makes PPARγ modulators attractive tools for PSDE control.

PLA2G7 is one of the highest-risk HCPs for PS degradation due to its strong activity against PS and tendency to persist through purification.^10,58^ Thus, even a modest decrease in the PLA2G7 level can potentially lead to meaningful improvement in PS stability and shelf life. For comparison, a previous study demonstrated that similarly modest (< 50 %) decrease in the cathepsin D level was sufficient to prevent mAb fragmentation.^59^ In a manufacturing setting, even a modest 5–10 % extension in product shelf life resulting from the reduction and control of HCPs would translate to millions of dollars of cost saving. Efforts to extend product shelf life may hopefully reduce the drug cost and improve the equitable access to life-saving medicines.

## Methods

### Cell line and cell culture operations

Two clones of a glutamine synthetase (GS) knockout CHO cell line, each carrying a monoclonal antibody (IgG mAb) (Clones A and B), were used for experiments. The clones were passaged in shake flasks in a humidified Multitron Cell incubator at 37 °C, 5 % CO_2_, and 140 rpm using a chemically defined CHO medium containing 12.5 µM methionine sulfoximine (MSX, Sigma-Aldrich). Exponentially growing cells were inoculated at 0.5 × 10^6^ cells/mL in Dynamis® culture media (Gibco) in Thompson optimum growth flasks with three biological replicates. Bioreactor cultures were maintained at the same conditions with the addition of two different feeds (Feed A and Feed B) every alternate day starting from Day 3 at 5.12 % *v*/*wv* (Feed A) and 0.60 % *v*/*wv* (Feed B). Glucose was spiked up to 6 g/L whenever the measured glucose level fell below 3 g/L. pH was adjusted to 7.2 with NaOH whenever the pH fell below 6.9. For Batch culture, Clone A was cultured for 7 days, in a 24-well plate in the same incubator at 300 rpm agitation speed. Glucose was supplied whenever the level fell below 3 g/L. SR1664 was added on day 0 and day 2 at 1 µM concentration. For the bioreactor run in the presence of small molecules, Clone A was batched in AMBR250 bioreactors (Sartorius Stedim) equipped with two pitched blade impellers and an open pipe sparger, seeded at 0.5 × 10^6^ cells/mL Dynamis® culture media (total volume: 190 mL). Feed A and Feed B were supplied every day starting from Day 3 at concentrations of 2.56 % *v*/*wv* and 0.30 % *v*/*wv*, respectively. The pH range and glucose targets were the same as in the shake flask run. Antifoam (0.5 % simethicone) was supplemented as needed, and sodium carbonate was added to control the pH. Dissolved oxygen was maintained at around 35 %. SR1664 was added to maintain a concentration of 1 × 10^-15^ mol/cell up to Day 6.

### Cell culture sample analysis

Daily cell culture samples were collected and immediately analyzed. Viable cell density and viability were assessed using the trypan blue exclusion method with a Cedex HiRes cell counter (Roche Diagnostics). pH, pO_2_, and pCO_2_, glucose, lactate, glutamine, and glutamate levels were determined using a Flex analyzer (Roche Diagnostics). Cell culture fluids were filtered through 0.22 µm PVDF membrane for long term storage and downstream analysis. Antibody titers (g/L) were measured using a HPLC based Protein A affinity column, specifically designed for monoclonal antibody analysis. Total lipase activity was determined with our proprietary assay using a fluorogenic substrate that structurally mimic a lipid.

### Protein A affinity chromatography

Protein A affinity chromatography was performed using MabSelect SuRe (Cytiva) resin packed into a column of 6.74 mL in volume with 19.7 cm bed height. The residence time for these experiments was 4 min. An ÄKTA™ avant 25 system was used to run these experiments. Approximately 30 g/L of HCCF was loaded onto the column on an AVANT system for each run. Before each run, the column was sanitized with 5 column volumes (CVs) of sanitization buffer (0.1 M NaOH). It was then equilibrated with 5 CVs of equilibration buffer (10 mM NaPO_4_, pH 6.5). After the HCCF loading step, 3 CVs each of three washes were performed. Wash 1 and Wash 3 were performed with the equilibration buffer and Wash 2 buffer consisted of 10 mM NaPO_4_ and 0.5 M NaCl at pH 6.5. Then, the elution step was carried out with 5 CVs of elution buffer (20mM CH_3_COONa at pH 3.7), during which the protein A product (PAP) was collected. Elution was followed by a strip step with 3 CVs of strip buffer (100 mM CH_3_COOH).

### Proteomics sample preparation

#### Cell pellet samples

Cell pellets were processed with the SPEED method^60^. Approximately 0.25 × 10^6^ cells were washed twice with PBS (1 mL) and pelleted by centrifugation (4 °C, 800 rcf, 1 min). The cells were re-suspended in TFA (5 μL) and incubated at room temperature for 2 min. To this were added, 1.5 M tris base (62 μL), 1 M TCEP (1 μL), and 1 M acrylamide (2 μL), and this mixture was incubated at 95 °C for 5 min. The protein concentration was determined with the Nanodrop A280 assay (Thermo Fisher Scientific) and adjusted to 0.25 g/L with 50 mM tris-HCl pH 8.0. Digestion was performed at 25 °C for 20 h with trypsin (50:1, Promega) and LysC (500:1, Fujifilm-Wako). The digested peptides (400 ng) were loaded on an Evotip Pure Evotip (Evosep) according to the manufacturer’s instruction.

#### HCCF samples

HCCF was processed with the SP3 method^61^. HCCF (20 μL) was incubated with 4 M urea, 25 mM TCEP-HCl, and 50 mM acrylamide in 50 mM tris-HCl pH 8.0 (total volume: 50 μL) at 60 °C for 30 min. An aliquot (20 μL) was incubated with Sera-Mag Carboxylate-Modified Magnetic SpeedBead (1:1 mixture of E3 and E7, 10 μL, 200 μg; GE Healthcare) and acetonitrile (70 μL) at 20 °C for 10 min with agitation at 500 rpm. The beads were washed three times with 80 % EtOH (200 μL) and then incubated with 50 mM tris-HCl pH 8.0 (20 μL) containing 0.4 μg trypsin and 0.1 μg LysC at 37 °C for 18 h. The peptide concentration was determined with the Nanodrop A280 assay and/or the BCA assay (Thermo Scientific) and adjusted to 0.02 g/L with 0.1 % formic acid before loading 20 μL (400 ng) on an Evotip Pure (Evosep) according to the manufacturer’s instructions.

#### ABPP samples

HCCF (100 μL) was incubated with DMSO (1 μL) or 1 mM FP-biotin DMSO solution (1 μL; WuXi App Tec) at 25 °C for 1 h and subsequently filtered through Zeba Spin Desalting Column, &K MWCO, 100 μL (Thermo Fisher Scientific). The protein concentration was determined with the Nanodrop A280 assay (Thermo Fisher Scientific). An aliquot (0.5 mg) was diluted to 0.5 g/L with 0.2 % SDS in PBS pH 7.1 and incubated with Pierce Streptavidin Magnetic Beads (20 μL; Thermo Fisher Scientific) at 25 °C for 1 h. Then, the beads were washed three times with 0.2 % SDS / PBS pH 7.1 (1 mL) and three times with 1 M urea / water (1 mL). On-bead digestion was performed in 50 mM tris-HCl pH 8.0 (50 μL) containing 0.1 μg trypsin (Promega) and 0.01 μg LysC (Fujifilm-Wako) at 25 °C for 20 h. The supernatant (20 μL) was loaded on an Evotip Pure (Evosep) according to the manufacturer’s instruction.

### Proteomics data acquisition

#### DIA–based quantitative proteomics

All samples were run on the Evosep One LC–Bruker timsTOF Pro 2 system with an Pepsep C18 column on the diaPASEF mode^62^. Peptides were separated on the Evosep One LC system with a Pepsep C18 column (15 cm × 150 μm, 1.9 μm) using the 30 samples per day method (mobile phase A: 0.1 % formic acid in water; mobile phase B: 0.1 % formic acid in acetonitrile; gradient time: 44.0 min; cycle time: 48.0 min; flow rate: 0.5 μL/min) (cell pellet) or Pepsep C18 column (8 cm × 100 μm, 3 μm) using the 100 samples per day method (mobile phase A: 0.1 % formic acid in water; mobile phase B: 0.1 % formic acid in acetonitrile; gradient time: 11.5 min; cycle time: 14.4 min; flow rate: 1.5 μL/min) (HCCF) and analysed on the Bruker timsTOF Pro 2 system in the diaPASEF mode. The diaPASEF method set both TIMS accumulation and ramp times at 100 ms and covered the ion mobility range 0.7–1.4 1/*K*_0_ and the mass range 300–1200 *m*/*z* with 25 varying-sized windows (cycle time: 2.76 s) (cell pellet) or 16 varying-sized windows (cycle time: 0.95 s). The variable window schemes were previously optimized with the py_diAID software^63^.

Spectral library was generated from repeated measurements of a sample pool, which was either a mixture of aliquots from all cell pellet samples or a mixture of aliquots from all HCCF samples, via the ion mobility–coupled gas phase fractionation (IM– GPF) approach. Peptides were separated on the Evosep One LC system with a Pepsep C18 column (15 cm × 150 μm, 1.9 μm) using the 15 samples per day method (mobile phase A: 0.1 % formic acid in water; mobile phase B: 0.1 % formic acid in acetonitrile; gradient time: 88.0 min; cycle time: 92.0 min; flow rate: 220 nL/min) (cell pellet) or Pepsep C18 column (8 cm × 100 μm, 3 μm) using the 60 samples per day method (mobile phase A: 0.1 % formic acid in water; mobile phase B: 0.1 % formic acid in acetonitrile; gradient time: 21.0 min; cycle time: 24.0 min; flow rate: 1.0 μL/min) (HCCF) and analysed on the Bruker timsTOF Pro 2 system in the diaPASEF mode. The IM–GPF method set both TIMS accumulation and ramp times at 100 ms and covered the ion mobility range 0.7–1.4 1/*K*_0_ and the mass range 300–1200 *m*/*z* with 100 effective 1-*m*/*z* windows (width: 2 *m*/*z*; overlap: 1 *m*/*z*) over 9 injections (cell pellet) or 50 effective 1-*m*/*z* windows (width: 2 *m*/*z*; overlap: 1 *m*/*z*) over 18 injections (HCCF).

#### ABPP

All samples were run on the Evosep One LC–Bruker timsTOF Pro 2 system with an Pepsep C18 column. The 30 sample per day method and the standard DDA PASEF method^64^ were used without modification.

### Proteomics data analysis

DDA PASEF data were analysed with MaxQuant^65^ v2.4.10. Use of .NET Core was disabled. Match between runs (MBR) was enabled. All other settings were left as default. All diaPASEF data were analysed with DIA-NN^66^ v1.8.1. Mass accuracies were set to 10 ppm for spectral library generation and to 15 ppm for all other processing. MBR was disabled. For the generation of the in-silico spectral library from sequence databases, the precursor charge range and the mass range were restricted to 2−4 and 300–1200 Th, respectively. The precursor and protein FDR thresholds were set to 1 % for all analyses. An in-house sequence database covering the proprietary product, the CHO proteome, and common contaminants was used in all analysis.

#### DIA–based quantitative proteomics

Precursors filtered by requiring identification in ≥ 80 % of sample pool runs, coefficient of variation across sample pool runs < 0.5, and global precursor and global protein Q values < 0.01. These precursors were then used to calculate protein quantities based on the Precursor.Normalised column using the MaxLFQ algorithm as implemented in the iq R package^67^. Missing values were imputed with a mixed imputation strategy: Protein quantities that were missing in < 5 % of the samples were imputed with the QRILC algorithm as implemented in the imputeLCMD R package; Values that were missing in ≥ 5 % of the samples were imputed with the k-nearest neighbour averaging algorithm as implemented in the same R package. Differential abundance analysis was performed using the limma R package^68^. The linear models were fitted protein group–wise using the lmFit function. The arrayWeights function^69^ was used in the analysis of the cell pellet samples. Clones or conditions were compared at each timepoint using the makeContrasts and contrasts.fit functions. The moderated t-statistics were computed using the eBayes function, allowing an intensity trend in the prior variance. The robust option in the eBayes function was enabled in the analysis of the cell pellet samples. Adjusted p-values were extracted using the topTable function. The Benjamini-Hochberg procedure^70^ was used for multiple testing. The results of differential expression analysis were used for GSEA^71^ (human ortholog genes were retrieved from Ensembl). GSEA was performed using the gseGO function from the clusterProfiler R package^72^ with following settings changed from the default values: nPermSimple = 10^5^; eps = 0. Gene Ontology was retrieved from the org.Hs.eg.db R package. Gene set over-representation analysis (Fisher’s exact test) was performed using the phyper function from the stats R package. A list of PPARγ target genes were retrieved from the PPARgene database^47^ and used as the gene set. All quantified genes were used as background. Hierarchical clustering was performed using the hclust function from the stats R package. The distance was computed as 1 – r (Pearson’s correlation coefficient) and the complete linkage method was used.

#### ABPP

Proteins were considered only if quantified in ≥ 50 % of the samples. Missing values were imputed with the QRILC algorithm as implemented in the imputeLCMD R package. Differential abundance analysis was performed using the limma R package^68^. The linear models were fitted protein group–wise using the lmFit function. The arrayWeights function^69^ was used. Conditions were compared at each timepoint using the makeContrasts and contrasts.fit functions. The moderated t-statistics were computed using the treat function^73^, testing significance relative to the two-fold change threshold. The trend and robust options were enabled. Adjusted p-values were extracted using the topTreat function^73^. The Benjamini-Hochberg procedure^70^ was used for multiple testing.

### Transcriptomics data analysis

Gene-level counts were retrieved from our internal data repository.^21^ Counts were filtered by expression level, normalized using the trimmed mean of median method, and converted to CPM values, using the edgeR R package^74^. Differential expression analysis was performed using the limma R package^68^. The linear models were fitted gene-wise using the lmFit function. The arrayWeights function^69^ was used. Clones were compared at each timepoint using the makeContrasts and contrasts.fit functions. The moderated t-statistics were computed using the eBayes function, allowing an intensity trend in the prior variance. The robust option in the eBayes function was enabled. P-values were extracted using the topTable function. These timepoint-wise results were combined to estimate cumulative effect of gene expression by taking a cumulative sum of fold-changes. The p-values were combined using the Fisher’s method and adjusted for multiple testing using the the Benjamini-Hochberg procedure^70^. These combined results were used for GSEA^71^ and gene set over-representation analysis as above.

### mRNA–protein correlation analysis

Correlation between mRNA and protein levels for each gene was estimated for overlapping genes (n = 4953) between the transcriptomic and (cell pellet) proteomic data. Because these data were not obtained from the same samples, average values for each clone and timepoint (n = 6) were matched and used to calculate the Spearman’s correlation coefficient (ρ). The correlation analysis result was used for GSEA^71^ and gene set over-representation analysis as above.

### Characterization of PS-80 degradation by MAX-UPLC-ESI-HRMS

The Experiments were performed on a Waters Acquity UPLC Classic coupled to a timeLJofLJflight (TOF) mass spectrometer (Xevo G2-XS, Waters) that was equipped with an ESI source. An Oasis MAX Column (2.1 x 20 mm, 80Å, 30 µm, Waters) was used for mixed-mode chromatography. Initial conditions were set at 99% solvent A (0.1% Triflluoroacetic acid in water) and 1% solvent B (0.1% trifluoroacetic acid in acetonitrile) and held for 3 min. Solvent B was then increased to 20% at 3.1 min, to 25% in 7 min. to 80% at 10.1 min and held for 5 min, to 99% at 15.1 min and held for 4 min, to 1% at 19.1 min, and held for 3 min. The flow rate was 0.4 mL/min and column temperature was set at 50°C. 5 μL (equivalent to 1 μg of PS80) of sample were loaded. For the mass spectrometer, the following parameters were set for the ion source: capillary voltage 3.0 kV, sampling cone voltage 40 V, collision voltage 10 V, source temperature 150°C, and desolvation temperature 500°C, and cone gas flow rate at 30 L/h, desolvation gas flow rate at 850 L/h. ESI fullLJscan mass spectra were recorded over the m/z range of 120−4000 in resolution mode. Waters MassLynx 4.2 software was used for MS data acquisition. Waters raw data files were imported into Expressionist 18.0 to process without any prior conversion. The data analysis method included the following processes specified with any parameters that were changed from their default settings: m/z recalibration using background ions observed in the m/z range of the peaks of interest, retention time (RT) range restriction with an RT maximum of 20 minutes, chemical noise subtraction with chromatogram smoothing, an RT window of 99 scans, quantile of 50%, intensity thresholding of 10, an RT structure removal of 5 scans, and m/z structure removal of 3 points, peak detection with a summation of 1 scan, an RT peak splitting ratio of 30% and a smoothing of 5 scans; isotope clustering with RT tolerance of 0.05 min, m/z tolerance of 25 ppm; singleton peak filter.

## Acknowledgements

T.T. and A.S. thank Merck & Co., Inc., Rahway, NJ, USA Postdoctoral Program. The authors thank the inLJprocess analytical group for titer and product quality measurements support. The authors thank Divya Chandra, Alex Dow, Hillary A Schuessler, and Douglas Richardson for discussion and support.

## Author contributions

A.S. performed all cell culture experiments. G.P. performed protein A purification. A.S. and G.P. coordinated total lipase activity assay. J.W. and R.Y. performed polysorbate degradation assays. S.M. provided the transcriptomics data. S.M. and X.L. supervised the study. T.T. conceived the study, performed all proteomics experiments, analysed transcriptomics and proteomics data, and wrote the first draft with A.S., S.M., and X.L. All authors contributed to finalizing the manuscript and approved the final version.

## Data availability

Data for this article are unavailable for proprietary reasons.

---

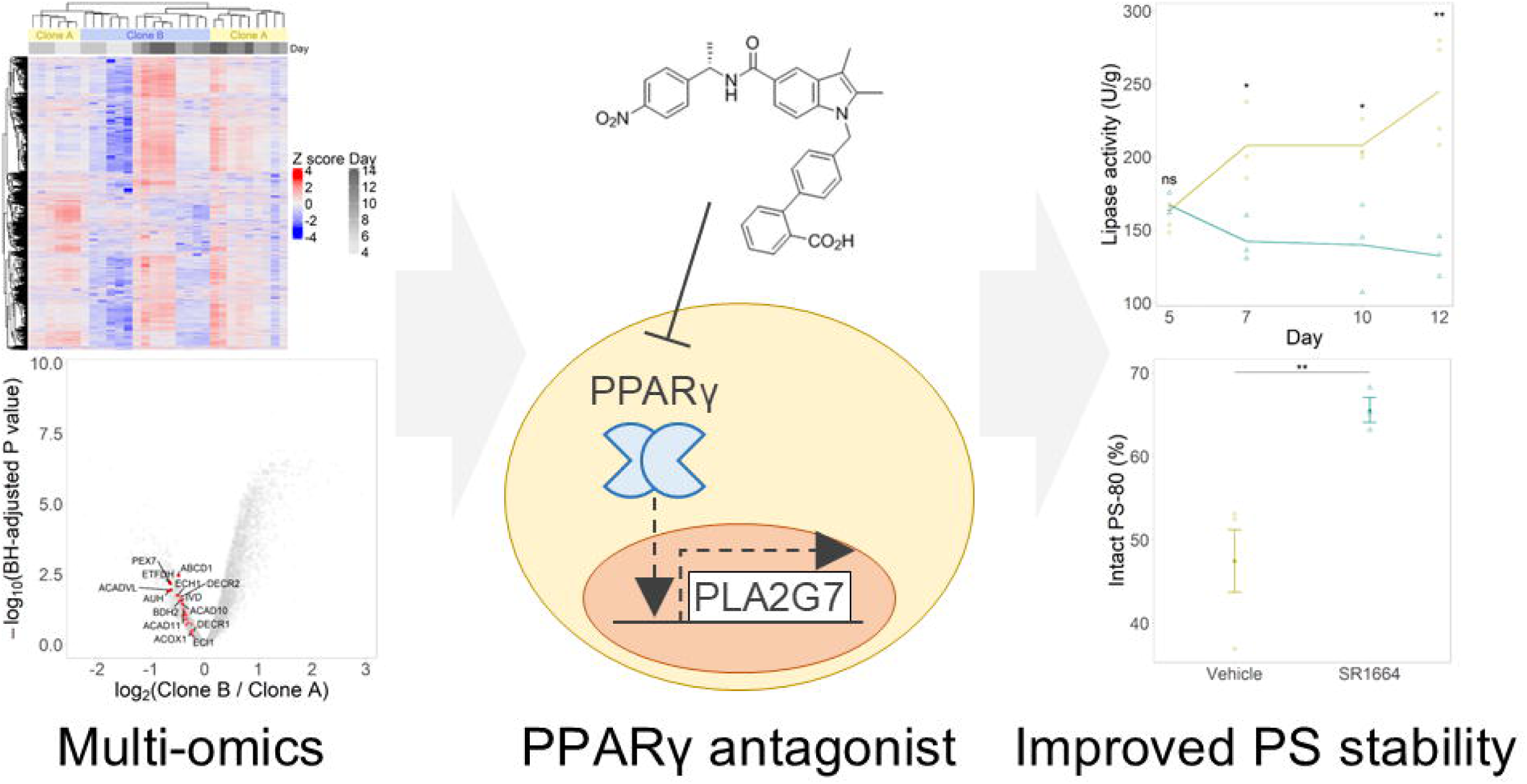

