## Supplementary Figures for "Discovery of Chemical Tools for Polysorbate-Degradative Enzyme Control in Biopharmaceutical Upstream Process via Multi-Omic Profiling of Host Cell Clones"

Supplementary Figure 1…………………………………………………………………………………………..S3

Supplementary Figure 2…………………………………………………………………………………………..S4

Supplementary Figure 3…………………………………………………………………………………………..S5

Supplementary Figure 4…………………………………………………………………………………………..S7

Supplementary Figure 5…………………………………………………………………………………………..S8

Supplementary Figure 6…………………………………………………………………………………………..S9

Supplementary Figure 7…………………………………………………………………………………………S10

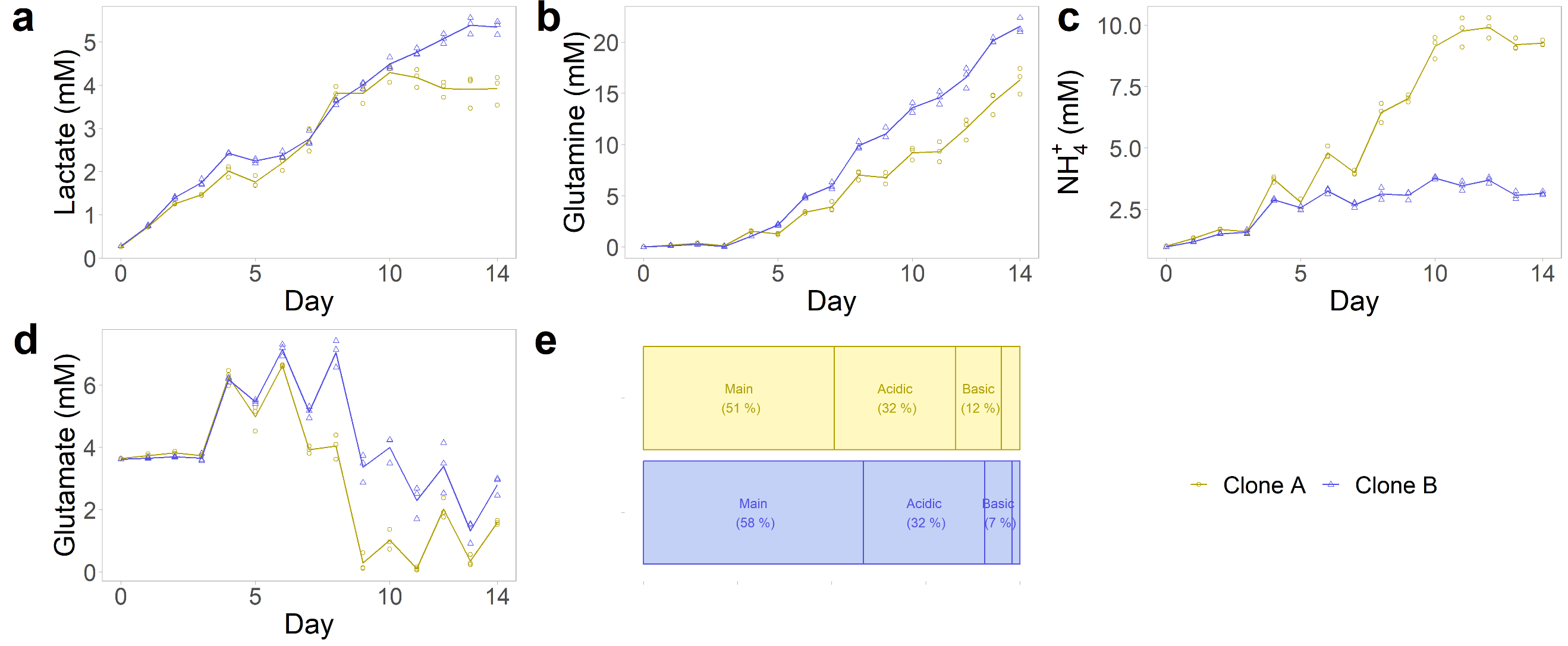
**Supplementary Fig. 1 | Fed-batch culture profiles.** **a** Lactate. **b** Glutamine. **c** Ammonium. **d** Glutamate. **e** Charge variant. n_Clone A_ = 3. n_Clone B_ = 3.

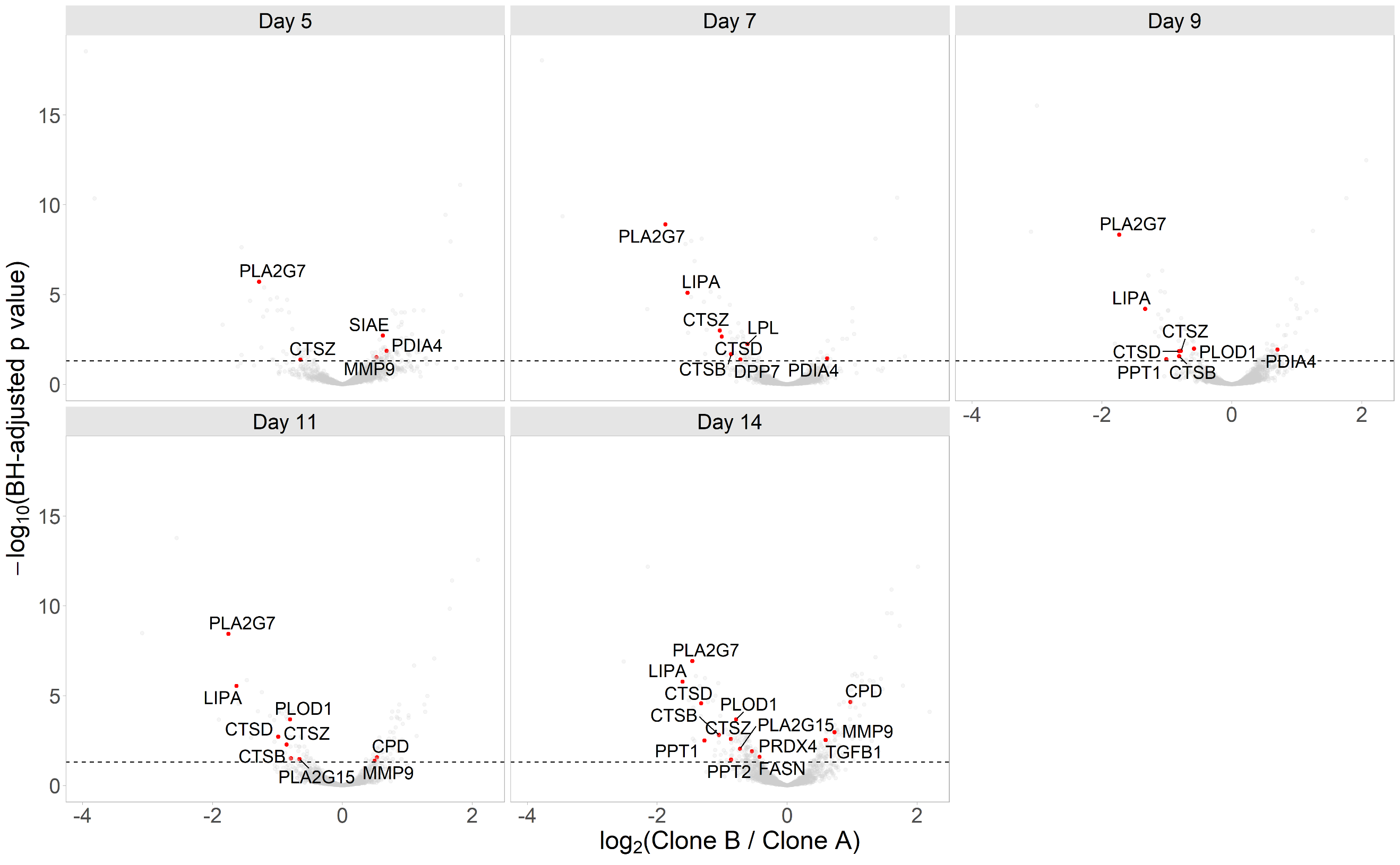
**Supplementary Fig. 2 | LC-MS/MS–based analysis of lipase and other high-risk HCP abundance in HCCF.** The horizontal line indicates adjusted p value < 0.05 in empirical Bayes–moderated *t*-test. Lipases and other high-risk HCPs with p < 0.05 are highlighted. n_Clone A_ = 3. n_Clone B_ = 3.

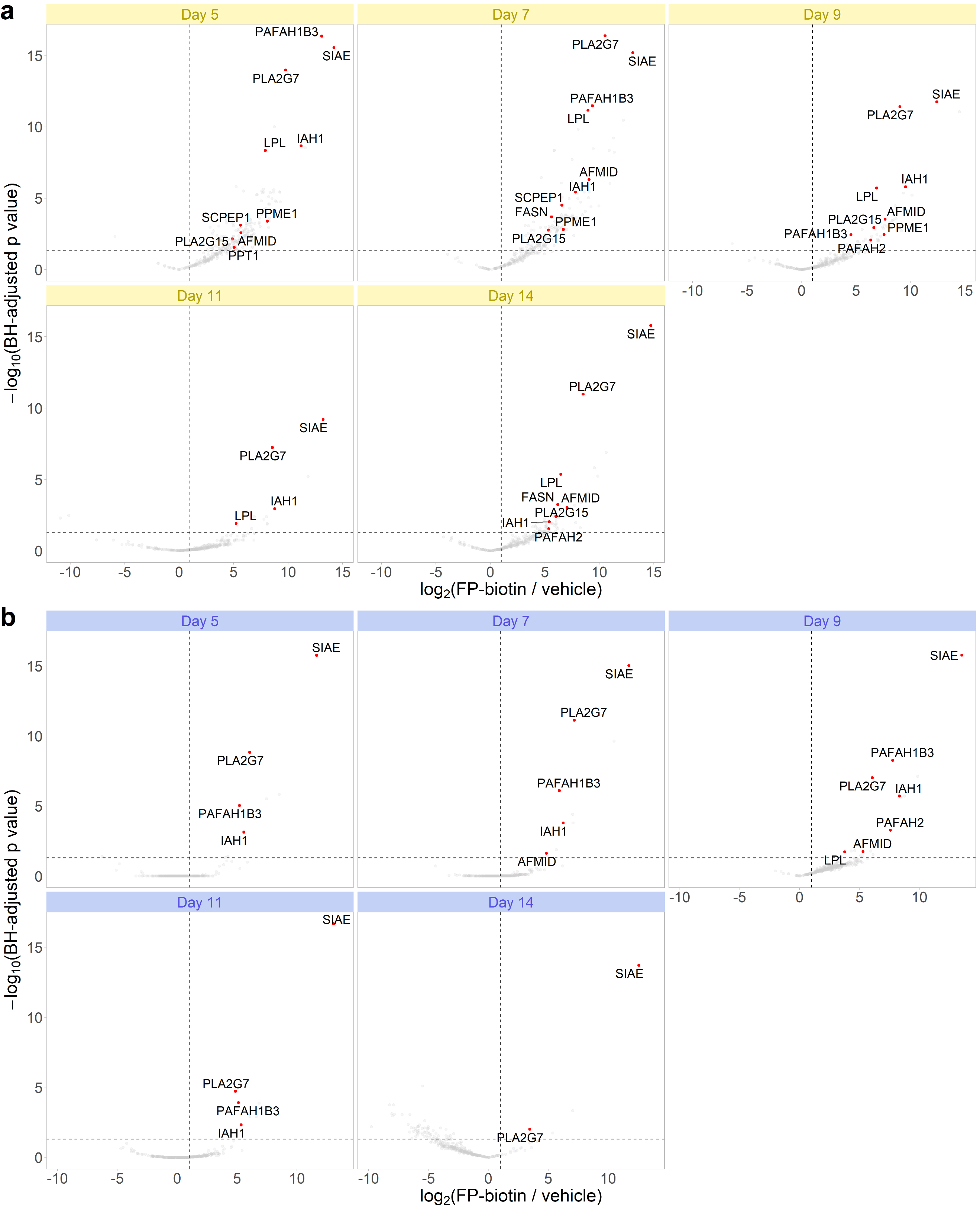
**Supplementary Fig. 3 | LC-MS/MS–based analysis of lipase activity in HCCP.** ABPP of lipases in Clone A (**a**) and in Clone B (**b**). The vertical line indicates log­_2_(FP-biotin / vehicle) > 1 and the horizontal line indicates adjusted p value < 0.05 in empirical Bayes–moderated *t*-test. Lipases with p < 0.05 are highlighted. n_FP-biotin_ = 3. n_vehicle_ = 3.

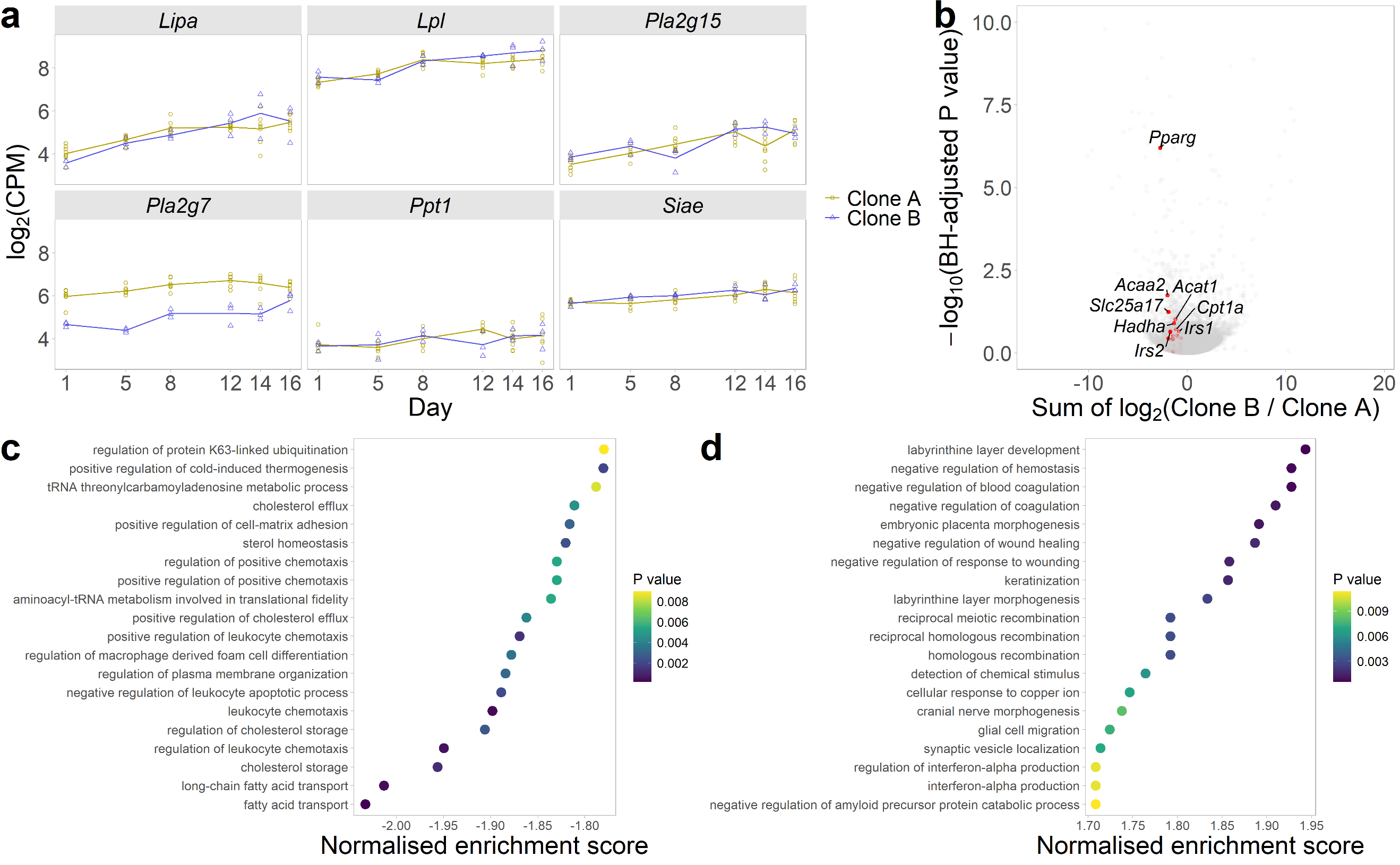

**Supplementary Fig. 4 | Transcript-level differential expression analysis. a** Time-course profiles of known PSDE transcripts. **b** Comparison of cumulative mRNA abundances between Clone A and Clone B at the eighth day with highlighted gene markers for fatty acid beta-oxidation leading-edge. Raw p values from timepoint-wise comparisons (empirical Bayes–moderated *t*-test) at the first, fifth and eighth days of culture were combined with Fisher’s method and corrected for multiple comparison with the Benjamini–Hochberg method. n_Clone A_ = 8. n_Clone B_ = 3. The plot was truncated at the adjusted p value of 1.0 × 10^-10^. GSEA of differentially expressed genes between Clone A and Clone B, showing top 20 pathways in Clone A (**c**) and top 20 pathways in Clone B (**d**).

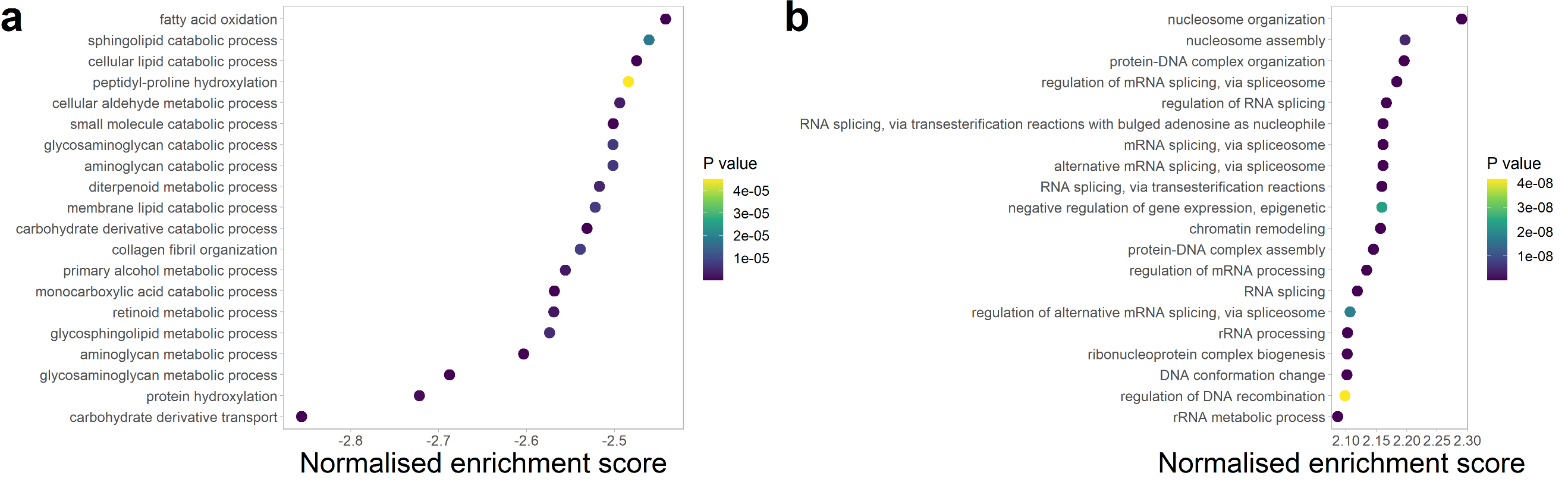
**Supplementary Fig. 5 | Protein-level pathway analysis.** GSEA of differentially abundant proteins between Clone A and Clone B, showing top 20 pathways in Clone A (**a**) and top 20 pathways in Clone B (**b**).

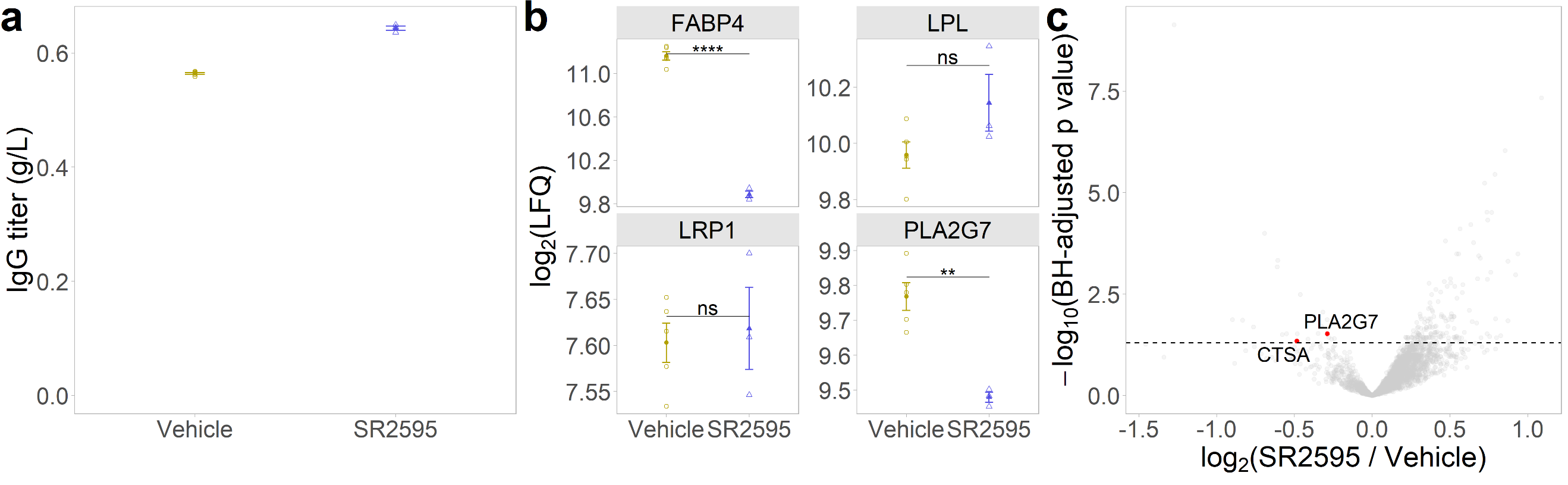
**Supplementary Fig. 6 | Selective reduction of PSDE levels in HCCF with PPARγ inverse agonist.** **a** SR2595 did not significantly affect IgG titer. **b** Protein levels of select PPARγ-target genes. Error bars represent the standard error of mean. Student’s *t*-test. ns: not significant; **: p < 0.01; ****: p < 0.0001. **c** Comparison of HCP levels between vehicle-control and SR2595-treatment groups, showing selective reduction of PLA2G7 and CTSA. The horizontal line indicates adjusted p value < 0.05 in empirical Bayes–moderated *t*-test. n_Vehicle_ = 6. n_SR2595_ = 3.

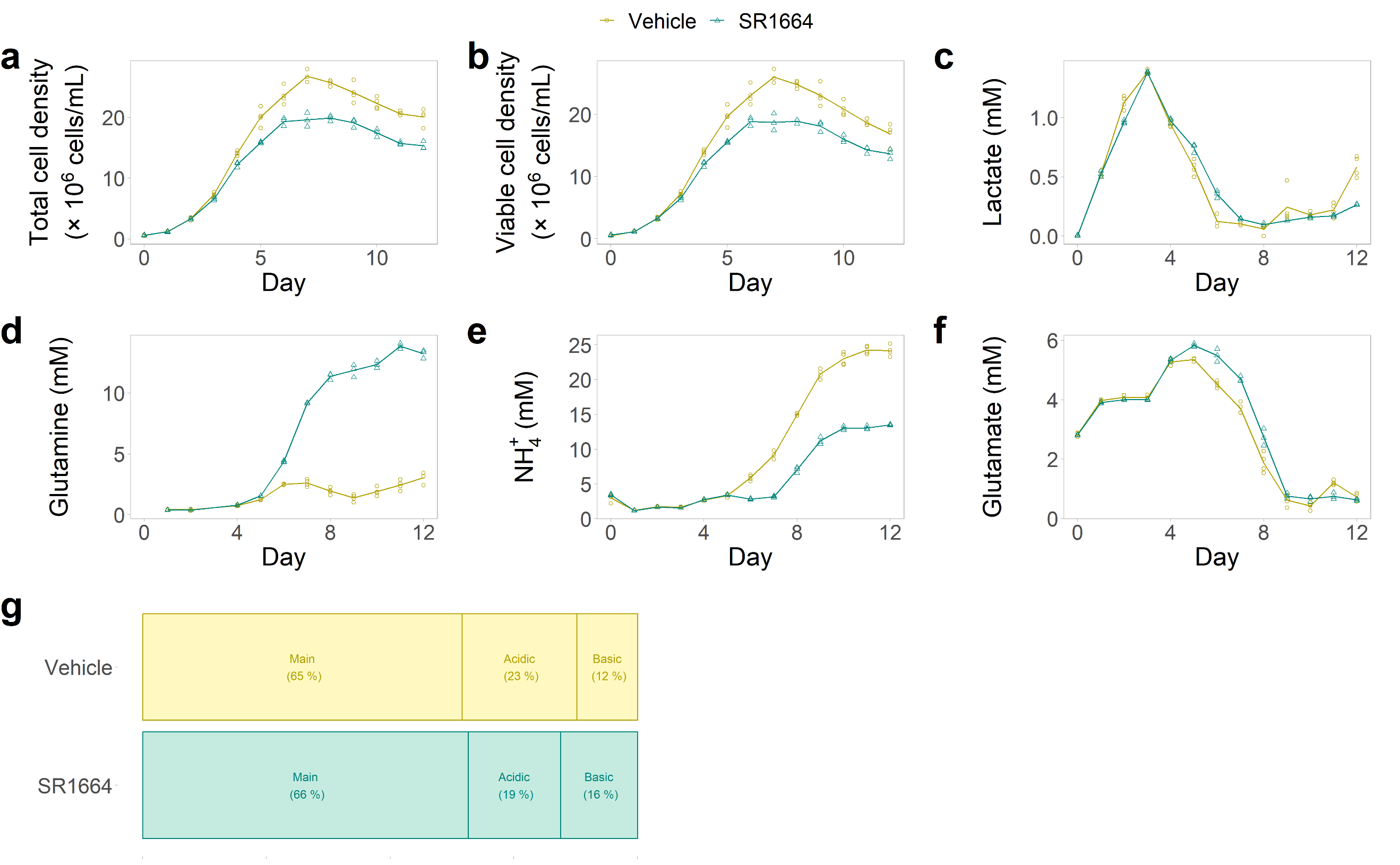
**Supplementary Fig. 7 | Culture profiles. a** Total cell density. **b** Viable cell density. **c** Lactate. **d** Glutamine. **e** Ammonium. **f** Glutamate. **g** Charge variant profile. n_Clone A_ = 4. n_Clone B_ = 3.
